# Neural microstates underlying categorical speech perception using Bayesian non-parametrics

**DOI:** 10.64898/2026.03.06.710122

**Authors:** Sultan Mahmud, Nahid Hasan, Kelsey Mankel, Mohammed Yeasin, Gavin M. Bidelman

**Affiliations:** Department of Electrical and Computer Engineering, University of Memphis, TN, USA; Center for Health System Improvement, College of Medicine, University of Tennessee Health Sciences Center, TN, USA; Department of Computer Science, Middle Tennessee State University, Murfreesboro, TN, USA; School of Communication Sciences & Disorders, University of Memphis, Memphis, TN, USA; Institute for Intelligent Systems, University of Memphis, Memphis, TN, USA; Department of Speech, Language and Hearing Sciences, Indiana University, Bloomington, IN, USA; Program in Neuroscience, Indiana University, Bloomington, IN, USA; Cognitive Science Program, Indiana University, Bloomington, IN, USA

**Keywords:** Auditory event-related potentials (ERPs), categorical perception, EEG microstates, Bayesian nonparametrics, brain–behavior relationships, machine learning, HDP-HMM

## Abstract

Categorical perception (CP) reflects the human auditory system’s ability to map continuous acoustic signals onto discrete categories. Understanding the relationship between neural dynamics and perceptual decisions is central to speech–language processing. Here, we implemented a data-driven approach using Bayesian nonparametrics and machine learning to characterize the relationship between auditory cortical responses and speech categorization behaviors. By applying a hierarchical Dirichlet process hidden Markov Model (HDP-HMM) to source-reconstructed event-related potential (ERP) data, we identified temporally distinct neural microstates that capture the evolving stages of speech categorization without imposing predefined temporal windows. Machine learning classifiers, including extreme gradient boosting (XGBoost), support vector machines, and random forests, were applied to decode prototypical (Tk1/5) versus ambiguous (Tk3) speech sound tokens from the microstate data. Using whole-brain activity, the XGBoost classifier achieved the highest decoding accuracy of 94.1% with an area under the curve (AUC) 94.1% within a specific encoding microstate occurring approximately 200-250 ms after stimulus onset. A reduced set of 15 informative brain regions identified via Shapley additive explanations (SHAP) yielded comparable classification performance (90.3% accuracy; AUC 90.0%), with many regions localized to frontal, temporal, and parietal regions in the left hemisphere. Furthermore, neural activity within these regions robustly predicted listeners’ behavioral identification slopes (*R*^2^ = 0.92, *p* < 0.00001), linking microstate-specific cortical dynamics to individual differences in perceptual gradiency of speech perception. These findings demonstrate that speech categorization emerges within temporally discrete neural microstates during early sensory–perceptual encoding and is supported by a selective, distributed cortical network with clear behavioral relevance.

## 1. INTRODUCTION

Categorical perception (CP) is a phenomenon that reflects the way individuals perceive and process speech sounds (Goldstone & Hendrickson, 2010; McMurray, 2022). Despite wide acoustic variability, CP allows listeners to extract, manipulate, and precisely identify sounds by grouping them into more manageable perceptual units (C. T. Miller & Cohen, 2010; E. K. Miller et al., 2003; Russ et al., 2007). CP of speech sounds emerges early in life but is also modified by language experience (Eimas et al., 1971). CP therefore plays a vital role in understanding the building blocks of speech perception and language processing.

Auditory event-related potentials (ERPs), recorded via scalp electroencephalography (EEG), provide temporally precise insights into the brain mechanisms of CP and speech processing (Celsis et al., 1999; Molfese et al., 2005). The N1-P2 components are voltage deflections that occur in auditory ERPs within the first 200 ms after sound presentation. The N1-P2 waves are associated with listeners’ speech identification <u>(Bidelman & Walker, 2017; Rizzi</u> <u>& Bidelman, 2024)</u>. Critically, larger or more differentiated N1-P2 responses have been shown to correspond to sharper behavioral identification boundaries and higher accuracy in distinguishing between speech categories. These findings are consistent with the notion that N1 and P2 are highly sensitive to speech processing and auditory object formation that is necessary to map sounds to meaning (Alain, 2007; Bidelman et al., 2013a; Wood et al., 1971a).

Machine learning (ML) classifiers have become increasingly important for decoding high-dimensional neuroimaging data, particularly in studies of speech perception where neural representations are distributed across space and time. Supervised ML models, such as support vector machines (SVM), random forests (RF), and extreme gradient boosting (XGBoost), offer powerful tools not only for capturing complex, multivariate relationships between neural signals and perceptual categories (Dutta et al., 2023; Hasan et al., 2024; Khamthung et al., 2024). For example, ML approaches suggest the human brain encodes the speech stimulus within *∼* 250 ms after stimulus onset, consistent with traditional ERP approaches (Masmoudi et al., 2012).

Previous studies by our group showed that the categorization of phonetic prototype vs. ambiguous speech sounds can be distinguished from ERP brain signals 180–320 ms after speech stimulus onset (Bidelman et al., 2020b; Mahmud, Ahmed, Al-Fahad, et al., 2020). This early period presumably reflects brain processing related to the sensory encoding of acoustic features in the speech signal. Meanwhile, speech identity can be decoded *≥* 300 ms after stimulus onset (Domenech & Dreher, 2010).

As decoding results are interpreted and incorporated into experimental design, ML-powered analyses become part of an iterative science cycle, guiding further hypothesis testing and deepening our mechanistic understanding of speech perception. Many machine learning approaches deliver strong predictive performance but are often criticized for their “black box” nature, which leaves neuroscientists and clinicians questioning how these models arrive at their decisions. For researchers in neuroscience, it is essential to move beyond simply achieving high accuracy to understand which neural features or brain regions are actually driving the model’s predictions. This motivation underpins the growing emphasis on interpretability: pinpointing which aspects of the brain’s activity are most important provides insight into the neurobiological foundations of cognition and behavior. To address this challenge, explainable artificial intelligence methods such as Shapley additive explanations (SHAP) enable principled identification of features that contribute most strongly to model predictions (Lundberg et al., 2019). In neuroimaging applications with high dimensional signals (e.g., multi-channel EEG), SHAP (Shapley Additive Explanations) facilitates (Lundberg et al., 2019; Salmi et al., 2025) the selection of informative brain regions while improving model generalization and linking decoding performance to neurobiological relevance.

Additionally, most EEG decoding studies are based on specific time windows that are determined *a priori* based on assumptions of the underlying time course of speech perception (Mahmud, Ahmed, Yeasin, et al., 2020; Mahmud et al., 2021c). While effective, such approaches may obscure the intrinsic temporal organization of neural activity underlying speech categorization. A more informative strategy is to allow the data itself to determine the number, timing, and duration of distinct neural states engaged during speech perception (i.e., categorizing speech tokens), without imposing predefined temporal boundaries. This data-driven approach offers a more informative strategy for revealing the brain’s natural segmentation of neural activity. Microstate-based modeling offers a principled framework for addressing this issue by segmenting continuous EEG activity into a sequence of quasi-stable neural states that capture recurring spatiotemporal patterns of brain activity (Fox et al., 2008; M. C. Hughes et al., 2015). Within this, the dwell time patterns—that is, the time spent in a particular microstate—can reveal important temporal aspects of neural dynamics which are often overlooked in static analyses of different “waves” of the ERPs (Fox et al., 2008; M. C. Hughes et al., 2015). These patterns help explain the spatiotemporal structure of neural activity, contributing to a deeper understanding of the underlying processes driving the model predictions (Vélez-Cruz, 2024).

The Hierarchical Dirichlet Process Hidden Markov Model (HDP-HMM) (Beal et al., 2001; Fox et al., 2011; Teh et al., 2004) offers an excellent Bayesian nonparametric framework for modeling and segmenting sequential data while inferring the number of latent states directly from the data. However, applying HDP-HMMs to large-scale neuroimaging datasets-such as high-density EEG, which may involve hundreds of channels and thousands of trials resulting in millions of data points—poses significant computational challenges. Conventional algorithms for inference in HMMs and HDP-HMMs are prone to local optima and often fail to adequately explore solutions with varying finite state spaces(M. C. Hughes et al., 2015). Stochastic optimization methods, although computationally efficient, typically assume a fixed number of states throughout training, which limits their adaptability and can lead to suboptimal performance in large or complex datasets (Foti et al., 2014; Johnson & Willsky, 2014). In contrast, Monte Carlo–based inference approaches allow state spaces to evolve and utilize the full dataset (Chang & Fisher III, 2014; Foti et al., 2014; Wang & Blei, 2012), but they require all sequences to be held in memory, resulting in substantial computational overhead and poor scalability. Consequently, prior neuroimaging studies have often adopted dimensionality-reduction strategies to improve tractability, including transforming multivariate EEG signals into representative univariate measures such as global field power (Michel & Koenig, 2018) or averaging data across trials or participants (Duc & Lee, 2019).

A promising strategy for overcoming these computational limitations is Memoized Variational Inference (moVB) (M. C. Hughes & Sudderth, 2013). Initially developed for Dirichlet Process (DP) mixture models and later extended to Hierarchical Dirichlet Process (HDP) topic models (M. Hughes et al., 2015), this approach was subsequently adapted for the sticky HDP-HMM to enable scalable inference on large datasets (M. C. Hughes et al., 2015). Unlike conventional inference methods that struggle with memory constraints or fixed state assumptions, moVB processes small subsets of sequences sequentially, allowing efficient learning without sacrificing model flexibility. This framework enables dynamic segmentation of multivariate EEG data across task-related and resting-state conditions while automatically inferring an appropriate number of latent microstates. Moreover, the incorporation of principled birth, merge, and delete operations prevents uncontrolled growth of the state space and promotes parsimonious solutions. Importantly, the memoized formulation supports incremental model updating as new observations become available, facilitating continuous learning and prediction of future data. Together, these properties make the sticky HDP-HMM with moVB particularly well suited for modeling the complex, high-dimensional neural dynamics underlying speech perception

Here, we aimed to apply state-of-the-art neural decoding techniques to determine how temporally distinct neural states in EEG/ERP activity support categorical speech perception. Our approach is fully data-driven and does not impose prior assumptions typically used in hypothesis-based analyses, such as predefined temporal windows, electrodes, or regions of interest that are assumed a priori to be discriminative (Al-Fahad et al., 2020; Mahmud, Ahmed, Yeasin, et al., 2020; Mahmud et al., 2021a).

To guide our investigation, we formulated the following hypotheses:

- Listeners with steeper identification slopes (i.e., stronger categorical perception) will exhibit distinct neural microstates with significantly different dwell times compared to listeners with shallower slopes.
- Specific microstate dwell times will be positively or negatively correlated with listeners’ behavioral gradience during speech identification, such that particular patterns of neural state expression underlie more (or less) categorical perception.
- Source-level microstate patterns can be linked to cortical regions implicated in speech categorization, helping to identify the neural generators associated with categorical processing.

A growing body of work has explored microstate-based analyses of EEG data to characterize speech categorization. For example, (Al-Fahad et al., 2021) applied Bayesian nonparametric microstate modeling to sensor-level EEG and demonstrated that distinct neural states reliably predicted listeners’ reaction-time decision speeds during speech categorization. That work established the utility of microstate-based decoding for linking neural dynamics to perceptual decision timing but was limited to scalp-level representations, which precluded inferences about underlying cortical generators.

The present study extends this prior work in several critical ways. First, rather than focusing on decision speed, we examine how neural microstates relate to individual differences in categorical perception strength, indexed by the gradiency (slope) of listeners’ identification functions. Second, we apply Bayesian nonparametric microstate modeling to source-reconstructed EEG, enabling identification of the cortical regions that contribute most strongly to speech categorization. To our knowledge, no previous study has examined how microstate dynamics derived from source-level EEG relate to behavioral gradience during categorical speech perception.

## 2. MATERIALS & METHODS

### 2.1 Participants

The data consisted of EEGs from N=49 young (15 males, 34 females; aged 18–33 years) adults that were recorded in our prior studies examining the neural basis of speech perception and auditory categorization (Bidelman et al., 2020b; Bidelman & Walker, 2017a; Mankel et al., 2020). All participants had normal hearing sensitivity (i.e., *<*25 dB HL between 500-4000 Hz). All but one listener was right-handed according to their Edinburgh Handedness scores (Oldfield, 1971) and had achieved a collegiate level of education. None reported any history of neurological disease. All participants were paid for their time and gave informed written consent in accordance with the declaration of Helsinki and a protocol approved by the Institutional Review Board at the University of Memphis.

### 2.2 Stimuli & task

We used a synthetic vowel continuum to assess the most discriminating spatiotemporal features while categorizing prototypical from ambiguous speech sounds (Bidelman et al., 2013a, 2014; Rizzi & Bidelman, 2024). Speech spectrograms are represented in Figure. 1A. Each token of the continuum was separated by equidistant steps acoustically based on the first formant frequency (F1) and perceived to categorically from “u” to “a”. Each speech token was 100 ms, including 10 ms rise/fall to minimize the spectral splatter in the stimuli. Tokens contained an identical voice fundamental frequency (F0), second (F2), and third formant (F3) frequencies (F0:150 Hz, F2: 1090 Hz, and F3:2350 Hz). To create a phonetic continuum that varied in percept from “u” to “a”, the F1 frequency was parameterized over five equal steps from 430 Hz to 730 Hz (Bidelman et al., 2013a).

**Figure 1:**
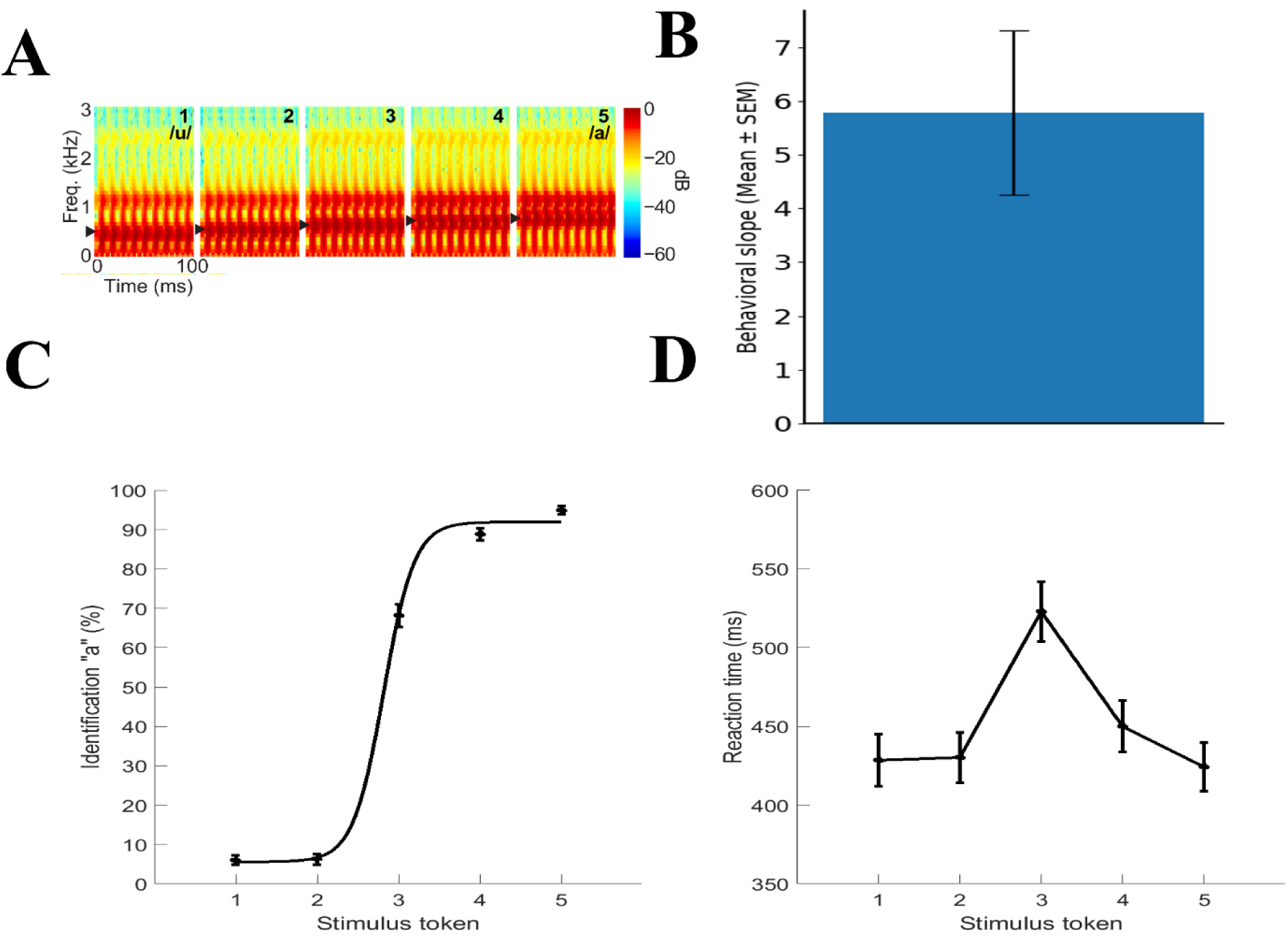
Speech stimuli and behavioral results. **A)** Acoustic spectrograms of the speech continuum from */u/* and */a/*. **B)** Behavioral slope. **C)** Psychometric functions showing % “a” identification of each token. Listeners’ perception abruptly shifts near the continuum midpoint, reflecting a flip in perceived phonetic category (i.e., “u” to “a”). **D)** Reaction time (RT) speeds for identifying each token. RTs are fastest for category prototypes (i.e., Tk1/5) and slow when classifying ambiguous tokens at the continuum midpoint (i.e., Tk3). Error bars = ±1 s.e.m.

Stimuli were presented binaurally at an intensity of 83 dB SPL through earphones (ER 2; Etymotic Research). Participants heard each token 150-200 times presented in random order. They were asked to label each sound in a binary identification task (“/u/” or “/a/”) as fast and accurately as possible. Their response and reaction time were logged. Behavioral identification functions revealed a sharp categorical boundary near the midpoint of the continuum (Tk3), as reflected in the psychometric functions and identification slopes (Figure.1B–C). Reaction times were longest for the category-ambiguous midpoint token and shorter for prototypical endpoint tokens (Figure.1D), consistent with increased decisional uncertainty during ambiguous speech categorization. The interstimulus interval (ISI) was jittered randomly between 400 and 600 ms (20 ms step and rectangular distribution) following listeners’ behavioral responses to avoid anticipating the next trial (Luck, 2005).

### 2.3 EEG recordings, behavioral measures, and data pre-processing procedures

During the behavioral task, EEG was recorded from 64 channels at standard 10-10 electrode locations on the scalp using an online reference located anterior to Cz (Oostenveld and Praamstra 2001). Continuous EEGs were digitized using Neuroscan SynAmps RT amplifiers at a sampling rate of 500 Hz. Subsequent preprocessing was conducted in the Curry 7 neuroimaging software suite, and customized routines coded in MATLAB. Ocular artifacts (e.g., eye-blinks) were corrected in the continuous EEG using principal component analysis (PCA) (Picton et al., 2000). The data were referenced to the common average reference and then filtered (1-100 Hz bandpass; notched filtered 60 Hz). Cleaned EEGs were then epoched into single trials (−200 to 800 ms, where *t* = 0 was stimulus onset).

### 2.4 EEG source localization

To disentangle the sources of CP-related EEG activity, we reconstructed the scalp-recorded responses by performing a distributed source analysis in the Brainstorm software package (Tadel et al., 2011). All analyses were performed on single-trial data. We used a realistic boundary element head model (BEM) volume conductor and standard low-resolution brain electromagnetic tomography (sLORETA) as the inverse solution within Brainstorm (Tadel et al., 2011). A BEM model has less spatial errors than other existing head models (e.g., concentric spherical head model). We used Brainstorm’s default parameter settings (signal to noise ratio [SNR]=3.00, regularization noise covariance = 0.1). From each single-trial sLORETA volume, we extracted the time-courses within 68 functional regions of interest (ROIs) across the left and right hemispheres defined by the Desikan-Killiany (DK) atlas (Desikan et al., 2006) (LH: 34 ROIs and RH: 34 ROIs). Single-trial data were then baseline corrected to the epoch’s pre-stimulus interval (−200-0 ms).

Since we aimed to decode prototypical (Tk1/5) from ambiguous speech (Tk3) stimulus presentations—a marker of categorical processing (Bidelman, 2015; Bidelman & Walker, 2019; Liebenthal et al., 2010)—we merged Tk1 and Tk5 responses since they reflect prototypical vowel categories (“u” vs. “a’). In contrast, Tk3 reflects a bistable percept—a category-ambiguous sound listeners sometimes label as “u” or “a” (Bidelman et al., 2020a; Bidelman & Walker, 2017a; Mankel et al., 2020). To ensure an equal number of trials and signal to noise ratio (SNR) for prototypical and ambiguous stimuli, we considered only 50% of the data from the merged (Tk1/5) samples.

### 2.5 Feature extraction

Previous empirical analysis demonstrated that averaging over 100 trials provided the best classification accuracy and maintained good SNR with computational efficiency (Al-Fahad et al., 2020; Mahmud, Ahmed, Al-Fahad, et al., 2020). We quantified source-level ERPs with a mean bootstrapping approach (James et al., 2013) by randomly averaging over 100 trials, 30 times with replacement, for each stimulus condition per participant (Al-Fahad et al., 2020).

### 2.6 Clustering Neural Data

Although the hierarchical Dirichlet process hidden Markov model (HDP-HMM) is a Bayesian nonparametric framework capable of inferring the number of latent states directly from the data, appropriate initialization is important for achieving stable and interpretable inference. We therefore performed an initial exploratory clustering analysis using Gaussian mixture models (GMMs) and the Bayesian Information Criterion (BIC) to estimate a plausible range of state complexity. This procedure was used solely to guide model initialization and did not constrain the HDP-HMM, which remained free to adapt the number of states through birth, merge, and deletion operations during inference.

To this end, we applied GMM clustering to the neural data using the expectation–maximization (EM) algorithm (Gupta & Chen, 2011). Compared with simpler methods such as k-means, GMMs provide greater robustness to noise and allow probabilistic cluster membership by modeling each cluster as a Gaussian distribution (Ramasso et al., 2022; Srivastava et al., 2023). We evaluated models with different covariance structures (spherical, tied, diagonal, and full) and varied the number of mixture components from 1 to 10. Model selection was guided by the BIC, which balances goodness of fit against model complexity. Across covariance types, the lowest BIC values consistently occurred at nine components when considering the full epoch (−200 to 800 ms). Accordingly, this value was used to initialize the emission structure of the HDP-HMM. BIC values across model configurations are shown in Figure 2.

**Figure 2:**
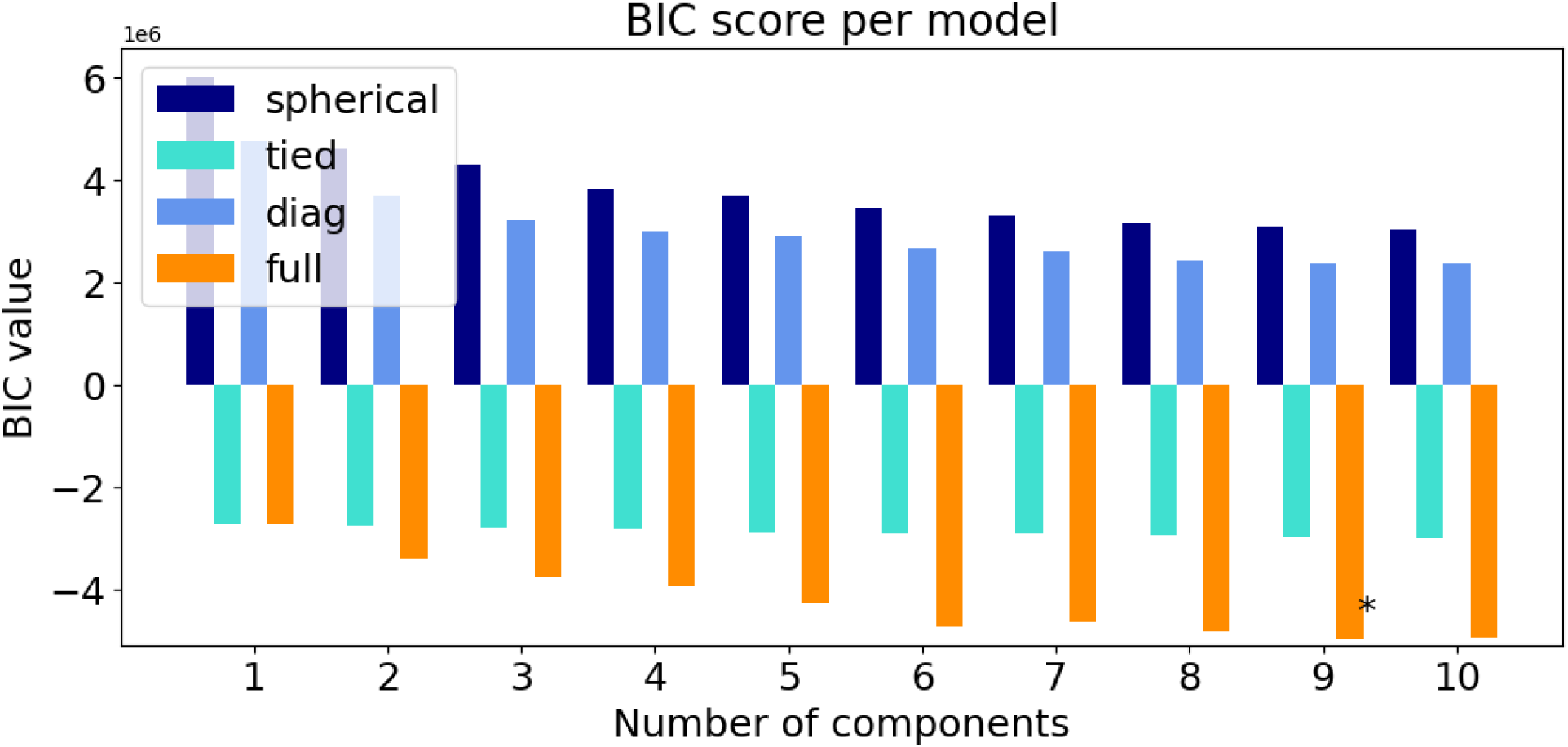
Clustering of ERP data while categorizing the prototypical (Tk1/5) vs. ambiguous (Tk3) speech sounds. Bayesian Information Criterion (BIC) scores with different co-variance methods and number of clusters. “*” represents the lowest BIC scores, and a suitable number of clusters to adequately described the data (here 9 clusters).

### 2.7 HDP-HMM

Our primary objective was to identify temporally distinct neural states that characterize brain dynamics during speech categorization without imposing prior assumptions about the timing or duration of processing stages. To this end, we employed the HDP-HMM (Fox et al., 2008) to model the temporal structure of the source constructed EEG data (Figure 3A) and infer sequences of latent neural states across the epoch.

**Figure 3.**
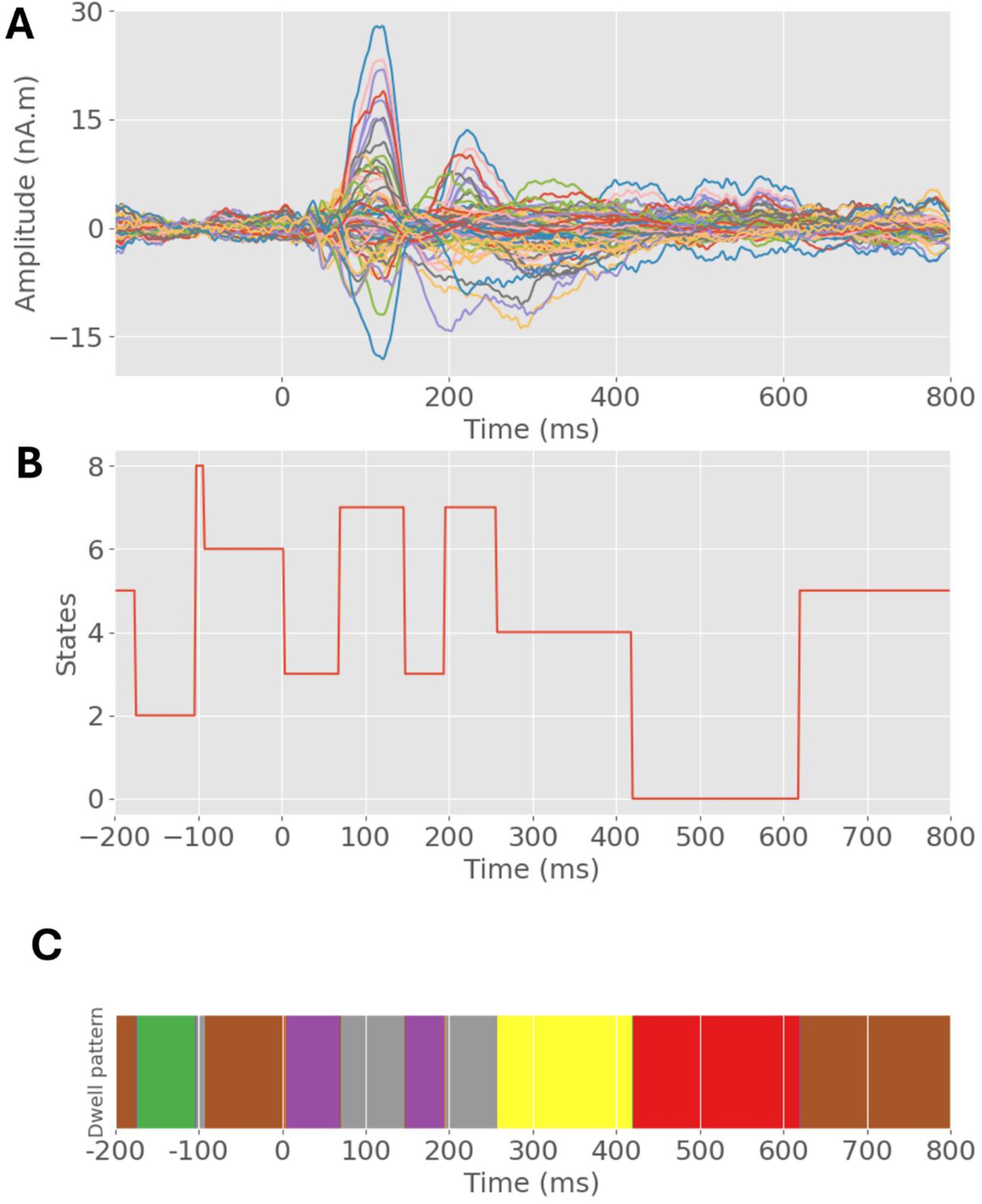
Representations of ERPs and dwell time patterns. **A)** Cortical ERP source signals derived from the 68 ROIs of the DK brain atlas. **B)** Dwell-time pattern showing the different microstates, and time within each state across the ∼1000 ms speech window t=0 represents the onset of the speech stimulus presentation. **C)** Corresponding state transitions of microstates corresponding to those shown in B (each color represents a distinct state). Brown: state 5; green: state 2; light gray: state 8; light brown: state 6; purple: state 3; gray: state 7; yellow: state 4; red: state 0.

Using the initialization described above, the HDP-HMM segmented the EEG time series into a sequence of microstates with variable durations, yielding both a state sequence (Figure 3B) and corresponding dwell-time patterns (Figure 3C). Dwell time reflects the duration for which the neural system remains in a given state and provides insight into the temporal organization of brain dynamics that is not captured by traditional fixed-window ERP analyses. Importantly, state boundaries and durations were determined directly from the data rather than imposed a priori. For each inferred microstate, we computed the mean ERP within each cortical ROI across the corresponding state-specific time window. These ERP mean features were then used as inputs to machine-learning classifiers to evaluate the discrimination of prototypical (Tk1/5) versus ambiguous (Tk3) speech tokens.

### 2.8 Classifiers (SVM, Random Forest, and XGBoost)

We used different genres of classical ML classifiers (e.g., SVM, RF, and XGBoost) to distinguish prototypical vowel vs. ambiguous speech tokens from the microstates neural data (e.g., mean ERP of microstate). These ML models learned to distinguish the data from the features (e.g., ERP) and corresponding class labels (e.g., prototypical speech [Tk1/5] and ambiguous speech token [Tk3]). Once the model was learned during the training phase, then features from unseen test data were used for prediction. We randomly shuffled and split the total data into training and test sets (80% and 20%, respectively) (Mahmud, Ahmed, Yeasin, et al., 2020). Classification performance (accuracy, area under the curve (AUC), and F1-score) was calculated by using standard formulas from the predicted and true class labels (Saito & Rehmsmeier, 2015). An excellent model has good separability and an AUC close to 1. On the other hand, a bad model has no separability with an AUC close to 0.

*SVM:* The SVM classifier learned the support vectors from the training data that comprise the feature attributes (e.g., source microstate ERP) and class labels (e.g., Tk1/5 and Tk3). The kernel function greatly affects SVM classifier performance, among other factors. The model hyperparameters (e.g., *C*, *γ*) reflect the classification performance (Hsu et al., 2003). During the training phase, we fine-tuned the model hyperparameters (e.g., *γ* and *C*) to detect the optimal values so that the model could accurately distinguish Tk1/5 vs. Tk3 speech tokens from the test data. A grid search approach was conducted to determine the optimal kernel, *γ*, and *C* values. We used five-folds cross-validation (Bhasin & Raghava, 2004) with kernels = ‘RBF’, 25 different values of (*C*, *γ*) in the following range for the *C* = [2*^-^*^1^ to 2^10^], and *γ* = [2*^-^*^4^ to 2^4^]. The optimal hyperplanes were fixed with maximum margin (i.e., maximum separation between the classes) and used for predicting the unseen test data.

*Random Forest (RF):* Random Forest is a robust ensemble learning algorithm that has gained significant attention in ML. In this work, we also used a parameter optimized RF classifier (Pal, 2005). The values of *n*_*estimators* and *max*_*depth* play an important role in the performance of the RF classifier (Pal, 2005). During the training phase, we conducted a grid search with the value range of *n*_*estimators* from 50 to 500 with step size 50; *max*_*depth* = [5, 10,40, 50]; *min*_*samples*_*split* = [2, 5, 10] to achieve maximum accuracy. In our data, we found optimal values of *n*_*estimators* = 250, *max*_*depth* = 40, *min*_*samples*_*split* = 2 provided the best classification accuracy.

*XGBoost:* We used a lightweight classifier called XGBoost with a base estimator decision tree classifier. The algorithm leverages a combination of regularization techniques, tree pruning, and parallel computing to enhance predictive accuracy and prevent overfitting. We conducted a grid search approach to achieve the optimal hyper-parameters with *learning*_*rate*: [0.05, 0.10]; *max*_*depth*: [3, 10, 15, 50, 100]; *min*_*child*_*weight*: [1, 3, 5, 10, 15, 20, 50]; *gamma*: [0.1, 0.2, 0.3, 0.4]; *colsample*_*bytree*: [0.3, 0.4, 0.7] The grid search approach showed the following optimal parameters: *learning*_*rate* = 0.10 *max*_*depth* = 10, *min*_*child*_*weight* = 5, *gamma* = 0.3, *colsample*_*bytree* = 0.3.

These three classifiers (SVM, RF, XGBoost) were trained on the same dataset and used to predict Tk1/5 vs. Tk3 using an unseen test dataset. The ERP attributes and corresponding class labels were submitted to the three classifiers in a similar manner. The classifiers’ performance metrics were calculated based on the predicted classes from the test data and the true class labels. We conducted a separate analysis of speech token classification using whole-brain data and data from the top 15 selected brain ROIs.

## 3. RESULTS

### 3.1 Classification of prototypical vowel vs. ambiguous speech tokens using whole-brain data in different microstates

To examine the best speech token classification accuracy across the different microstates, we applied different genres of classifiers (SVM, RF, and XGBoost) of the whole brain data (e.g., 68 brain regions). Classification accuracy results obtained by SVM, RF, and XGBoost in the different microstates are presented in the Figure. 4. All classifiers produced the highest accuracy in the 197-258 ms time window (SVM classifier accuracy: 92.32%, RF: 92.20%, and XGBoost: 94.10%). XGBoost showed the highest accuracy among all the classifiers and time windows (i.e., microstates); classification accuracies and other performance metrics for XGBoost specifically for the encoding time window (0 to 260 ms; microstates 3 and 7) are highlighted in Table 1. Notably, microstates 3 and 7 roughly correspond to time windows associated with the canonical auditory N1 and P2 waves reflecting the neural registration of sound in auditory cortex (Figure 3A). Our data are therefore consistent with previous work that demonstrates neural categorization of phonetic prototype vs. ambiguous speech sounds approximately 180–320 ms after stimulus onset using traditional ERP brain signals (Bidelman et al., 2020c; Mahmud, Ahmed, Yeasin, et al., 2020; Mahmud et al., 2021b).

**Figure 4:**
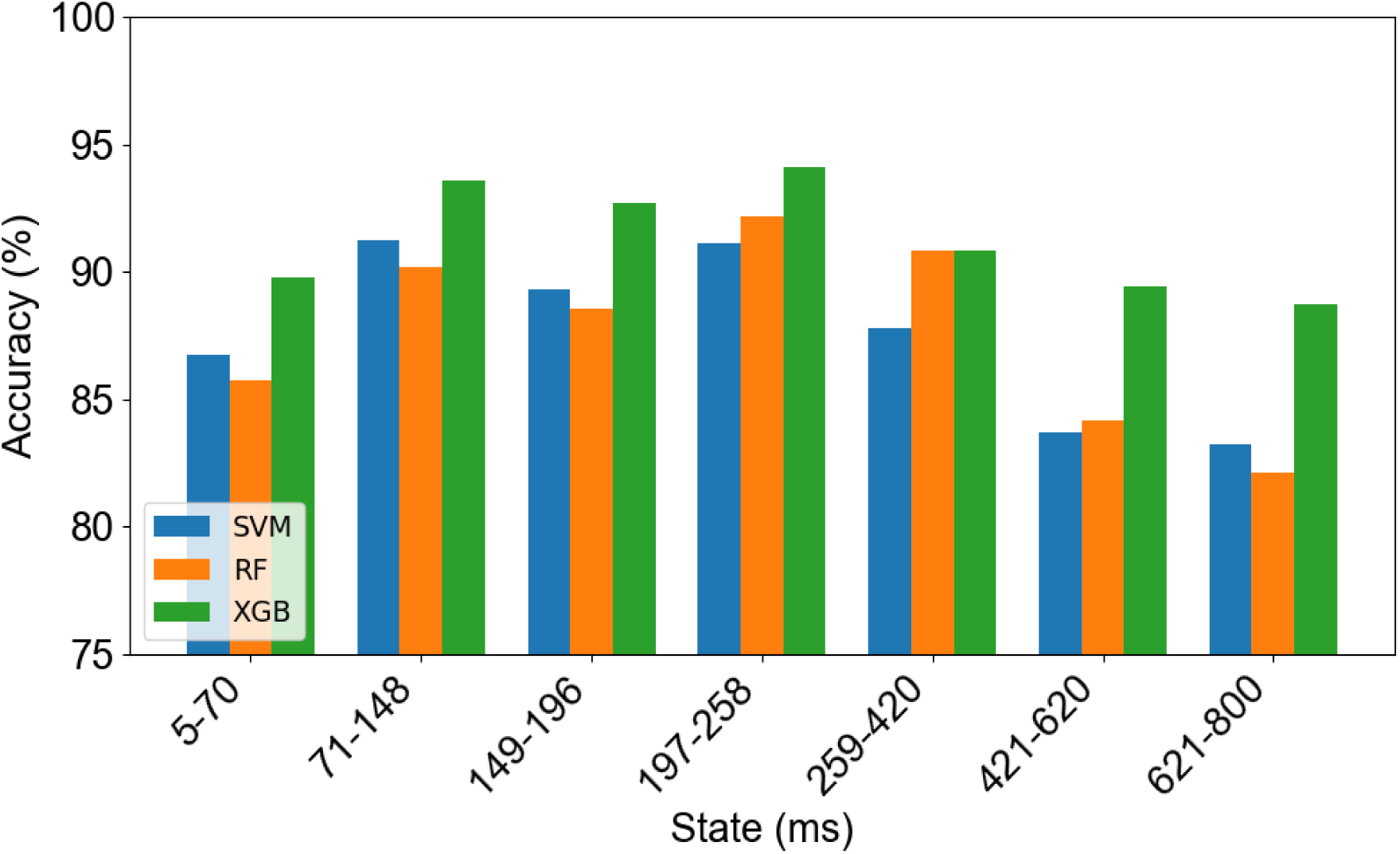
Prototypical speech vs. ambiguous speech token classification accuracy on different EEG microstates over the entire post-stimulus onset timeframe using SVM, RF, and XGBoost classifiers.

**Table 1:** XGBoost classifier performance metrics (%) during the encoding window (0–260 ms) for distinguishing prototypical vowel vs. ambiguous speech tokens in microstates 3 and 7.

| Microstate | Time Window (ms) | Average measure (%) | Whole-brain data | Top 15 ROIs |
| --- | --- | --- | --- | --- |
| 3 | 5–70 | Accuracy | 89.82 | 82.21 |
|  |  | AUC | 89.13 | 82.20 |
|  |  | Precision | 89.00 | 82.00 |
|  |  | Recall | 89.00 | 82.00 |
|  |  | F1-score | 89.00 | 82.00 |
| 7 | 71–148 | Accuracy | 93.61 | 87.82 |
|  |  | AUC | 93.11 | 87.27 |
|  |  | Precision | 93.00 | 88.00 |
|  |  | Recall | 93.00 | 88.00 |
|  |  | F1-score | 93.00 | 88.00 |
| 3 | 149–196 | Accuracy | 92.27 | 89.15 |
|  |  | AUC | 92.10 | 89.20 |
|  |  | Precision | 93.00 | 89.00 |
|  |  | Recall | 92.00 | 89.00 |
|  |  | F1-score | 92.00 | 89.00 |
| 7 | 197–258 | Accuracy | 94.12 | 90.28 |
|  |  | AUC | 94.13 | 90.17 |
|  |  | Precision | 94.00 | 90.00 |
|  |  | Recall | 94.00 | 90.00 |
|  |  | F1-score | 94.00 | 90.00 |

### 3.2 Classification of prototypical speech vs. ambiguous speech token using top 15 ROIs’ data

To assess feature importance and guide dimensionality reduction, we used SHAP (Lundberg et al., 2019) to quantify the contribution of each ROI to classification performance. Based on SHAP-based feature attribution, we identified a reduced subset of 15 ROIs with the highest average absolute SHAP values for subsequent analyses. This subset demonstrated strong generalization capability and robust classification performance in distinguishing prototypical (Tk1/5) from ambiguous (Tk3) speech tokens. The top-ranked ROIs are shown in Figure 5. These ROIs demonstrated strong generalization capability and robust classification performance in distinguishing prototypical (Tk1/5) from ambiguous (Tk3) speech tokens from the EEG. Using ERPs extracted from these top-ranked ROIs, we classified Tk1/5 versus Tk3 within the speech encoding interval, corresponding to microstates 3 and 7 (0-260 ms). Classification performance for the best performing classifier, XGBoost, is reported in Table 1. Noticeably, the time window corresponding to 197-258 ms showed the highest classification ability for Tk1/5 vs. Tk3 (accuracy 90.28%, with AUC 90.17% and F1-score, precision, and recall 90.00%).

**Figure 5:**
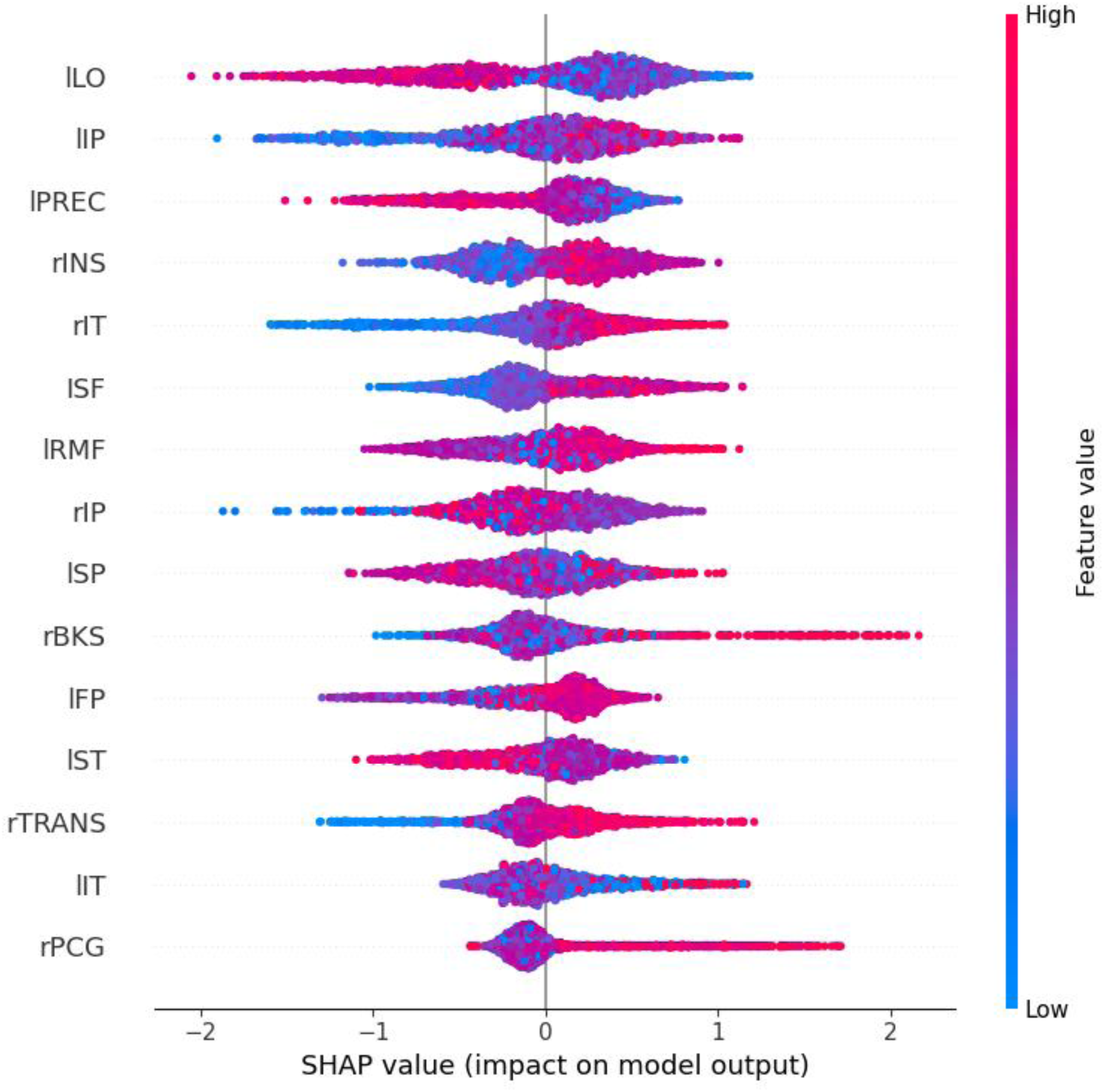
Density scatter plot of SHAP values obtained from the XGBoost classifier. Each point reflects the contribution of a single observation for a given ROI to the XGBoost model prediction, quantified by its SHAP value. Positive values favor one class, negative values favor the other. l/r: left/right; LO: lateral occipital; IP: inferior parietal; PREC: precuneus; INS: insula; IT: inferior temporal; SF: superior frontal; RMF: rostral middle frontal; SP: superior parietal; BKS: banks of the superior temporal sulcus; FP: frontal pole, ST: superior temporal; TRANS: transverse temporal; PCG: posterior cingulate.

### 3.3 Brain-behavior correspondences

Behavioral identification (%) functions and reaction time (ms) for speech categorization are shown in Figure. 1C and Figure. 1D, respectively. Listeners responses abruptly shifted in speech identity (/u/ vs. /a/) near the midpoint of the continuum, reflecting a change in perceived category. The behavioral speed of speech labeling [i.e., reaction time (RT)] were computed as listeners’ median response latency for a given condition across all trials. RTs outside of 250-2500 ms were deemed outliers and excluded from further analysis (Bidelman et al., 2013b; Bidelman & Walker, 2017b). Listeners spent more time classifying the ambiguous (Tk3) than prototypical speech tokens (e.g., Tk1/5), consistent with categorical hearing (Pisoni & Tash, 1974). For each continuum, the identification scores were fit with a two parameters sigmoid function; 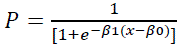, where *P* is the proportion of the trial identification as a function of a given vowel, *x* is the step number along the stimulus continuum, and *β0* and *β1* are the location and slope, respectively, of the logistic fit estimated using the nonlinear least-squares regression (Bidelman et al., 2014; Bidelman & Walker, 2017b). The mean slope of the sigmoidal psychometric function is presented in Figure 1B, reflecting the strength of listeners’ CP (i.e., steeper slopes correspond to stronger, more discrete vowel categorization).

To evaluate the behavioral relevance of the brain regions identified via SHAP, we conducted multivariate regression using weighted least squares (WLS) regression (Ruppert & Wand, 1994). Regression models were computed between the 15 ROIs identified in the decision interval and listeners’ behavioral slopes (Fig. 1B), which indexes their degree of categorical hearing (i.e., higher slope values indicate sharper distinction between phonetic categories). For each selected ROI, we extracted the mean ERP amplitude within the time window yielding the highest classification performance (197–258 ms; microstate 7), averaged across Tk1/5 and Tk3 conditions. Mean ERP values from the 15 ROIs were subsequently regressed simultaneously against listeners’ behavioral identification slopes. The inverse of the absolute error values of the ordinary least squares were used as weights in the WLS to reduce the effect of heteroscedasticity (Seabold & Perktold, 2010; *Weighted Regression in SAS, R, and Python*, n.d.). The multivariate model robustly predicted listeners’ behavioral CP from neural data (R^2^ = 0.918, *p*<0.00001; Table 2), demonstrating the selected 15 ROIs identified via SHAP carried behaviorally relevant information regarding the gradiency of listeners’ phonetic perception.

**Table 2:**
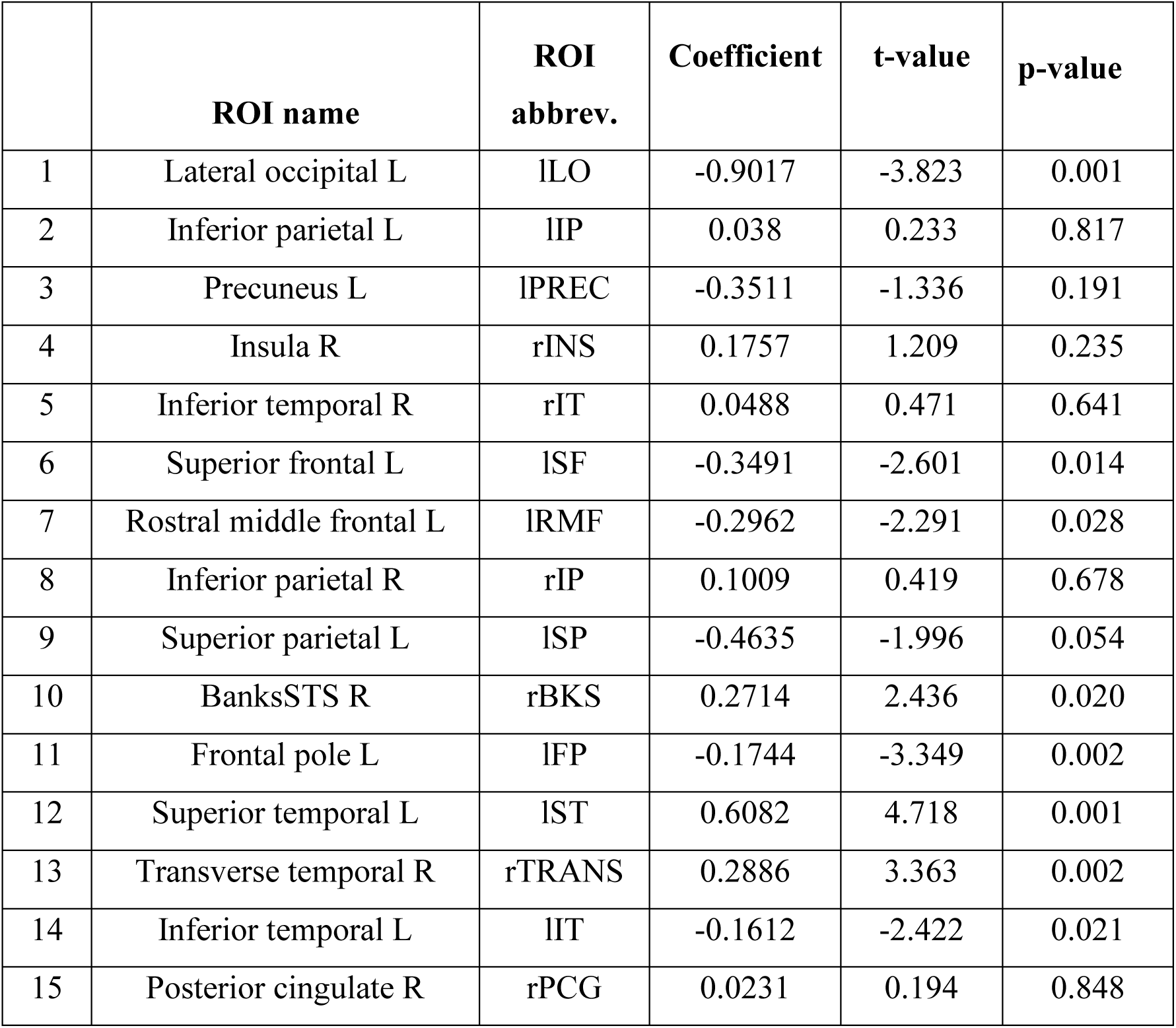
WLS regression results describing how individual brain ROIs predict behavioral CP.

## 4. DISCUSSION

In the current study, we combined data-driven microstate segmentation of EEG, ML-based neural decoding, and brain-behavior modeling to investigate how distributed cortical dynamics support categorical speech perception. This study represents a novel advance over previous hypothesis-driven analyses that relied on fixed sliding-window approaches (Mahmud, Ahmed, Yeasin, et al., 2020; Mahmud et al., 2021b). Our approach avoids arbitrary temporal segmentation of EEG/ERP responses by allowing the data itself to determine the number, timing, and duration of neural states involved in the perceptual process of speech categorization. Our results demonstrate that temporally distinct neural microstates arising during early-to-mid auditory evoked response latencies robustly distinguish prototypical from ambiguous speech tokens and account for substantial inter-individual variability in categorical perception, indexed by listeners’ identification slopes. Collectively, these findings provide converging evidence that categorical speech perception is grounded in early sensory–perceptual encoding processes from a distributed but relatively compact brain network and this information can be captured through assumption-free modeling of neural dynamics.

### 4.1 Microstate dynamics and neural decoding of speech categories

Using a sticky HDP-HMM, we identified distinct microstates in the post-stimulus epoch that distinguish neural responses to prototypical versus ambiguous speech sound tokens. While previous work has shown that categorical distinctions are strongest between ∼180–320 ms, such analyses typically relied on fixed sliding windows or predefined components (Mahmud, Ahmed, Yeasin, et al., 2020; Mahmud et al., 2021c) (Mahmud et al., 2021b) (Bidelman et al., 2020b; Mahmud, Ahmed, Al-Fahad, et al., 2020). Our microstate-based approach reveals that neural representations relevant to speech categorization are confined to specific, transient states whose boundaries vary dynamically over time. Across classifiers, the strongest decoding performance consistently emerged approximately 197–258 ms after stimulus onset. This time window overlaps with the auditory P2 component, which has been repeatedly linked to phonetic feature encoding, auditory object formation, and early categorical mapping of speech sounds (Alain, 2007; Bidelman et al., 2020a; Mahmud, Ahmed, Yeasin, et al., 2020; Masmoudi et al., 2012). We note that P2 is also a relatively high-amplitude evoked response and may therefore afford favorable signal-to-noise characteristics. However, the present findings do not rely on predefined temporal windows and instead identify this interval through unsupervised, data-driven state segmentation, suggesting that the observed decoding performance reflects temporally specific neural dynamics relevant to speech categorization rather than an a priori assumption about component structure. Importantly, these results extend prior ERP decoding studies by demonstrating that the temporal locus of categorical speech processing can be discovered directly from the data, rather than imposed by hypothesis-driven analyses.

The XGBoost classifier achieved higher classification accuracy than RF or SVM classifiers using both whole-brain data and a reduced set of 15 ROIs. Unlike RF, XGBoost includes regularization parameters and pruning features that reduce overfitting and enhance learning performance. Meanwhile, SVM can underperform with many features or when the decision boundaries are highly nonlinear. In several cases, XGBoost has been more effective than RF and/or SVM for classifying EEG data including seizure detection (Hong et al., 2022; Kode et al., 2024) sleep deprivation (Choi et al., 2018) or recognition of emotional states (Cowie et al., 2008). Though classification accuracy for all three models was within 10% for all microstate-defined time windows, XGBoost ultimately proved a better fit for our data, underscoring the importance of temporally adaptive feature extraction for capturing the neural basis of speech perception.

### 4.2 Reduced cortical representations preserve categorical information

A key finding of this study is that high classification accuracy was maintained even when neural decoding was restricted to a limited set of 15 cortical regions identified via SHAP-based feature attribution. Although classification performance using these regions was modestly lower than that with whole-brain data, the retained accuracy (∼90%) indicates that categorical speech information is concentrated within a distributed but selective cortical network. This suggests that not all cortical regions contribute equally to speech categorization and that meaningful dimensionality reduction can preserve behaviorally relevant information. The ability of a compact regional subset to support robust decoding further underscores the efficiency of cortical representations in describing a complex human behavior such as speech perception.

Most identified brain regions most relevant for speech token classification were located in the left hemisphere, consistent with the well-established left-lateralization of language processing (Eimas et al., 1971; Hull & Vaid, 2006; Tzourio et al., 1998)]. Regions within temporal and frontal cortices were especially prominent, supporting hierarchical models in which early acoustic representations in auditory cortex interact with higher-order linguistic and decision-related areas to form categorical percepts (Beach et al., 2021; Bonetti et al., 2024; Humphries et al., 2014; Yin et al., 2020; Zhou et al., 2022). The current results are also highly consistent with our previous findings indicating the importance of left hemisphere frontotemporal networks for encoding and decision-making processes during speech categorization (Mahmud et al., 2021b). Notably, engagement of left auditory cortex has been shown to play a critical role in shaping individual differences in the gradiency and categoricity of phonetic categorization, as indexed by identification slopes (Rizzi & Bidelman, 2024). These data-driven temporal segments highlight the importance of temporally precise feature extraction within auditory cortex for speech categorization. Additionally, the parietal lobe, particularly the inferior parietal lobe bilaterally, was also a strong contributor to classifying speech phonemes. These results support theories of dorsal and ventral streams within frontal, temporal, and parietal networks as important for speech-language processing (Hickok & Poeppel, 2000, 2004, 2007).

### 4.3 Brain–behavior coupling links microstates to perceptual outcomes

Regression analysis showed that neural responses across a subset of 15 brain ROIs between 197-258 ms explains a substantial proportion of variance in the slope of listeners’ behavioral identification. This indicates that roughly 50 ms of dynamic brain activity is necessary to explain the gradiency/categoricity in listeners’ phonetic perception. Traditional ERP measures during the timeframe of stimulus encoding (∼250 ms) suggest components like N1 and P2 reflect neural activity from several neural regions, primarily within the superior temporal gyrus (STG), auditory association areas, and even the reticular activating system (Crowley & Colrain, 2004; Näätänen & Picton, 1987; Woods, 1995). Evidence suggests the left STG is critical for phonetic feature extraction and phonological category encoding, and neural responses from this region are associated with steeper behavioral slopes (indicating sharper categorical boundaries) during speech categorization (Mahmud et al., 2021b; Ou & Yu, 2022; Rizzi & Bidelman, 2024; Russ et al., 2007; Yi et al., 2019). Similarly, engagement of the *right* transverse temporal gyrus (primary auditory cortex) positively predicted behavioral slope, highlighting the importance of high-fidelity acoustic encoding in the non-dominant hemisphere in shaping categorical decisions (Bhaya-Grossman & Chang, 2022; Wood et al., 1971b). Indeed, several key ROIs identified in our study were located in the temporal lobes, including specific subdivisions of STG. For example, the right transverse temporal lobe and bank of the STS contributed to speech categorization, consistent with their role in integrating acoustic and higher-order speech features, particularly under conditions of perceptual ambiguity (Mahmud et al., 2021c; Wood et al., 1971b). Together, these findings demonstrate that coordinated processing across primary, secondary, and integrative auditory regions is central to the precise mapping of speech sounds to category representations (Mahmud et al., 2021b).

Notably, areas in the frontal lobe also contributed to explaining perceptual responses. The superior frontal lobe is known for its role in working memory, attention, and other cognitive functions (Klingberg et al., 2002; Nyberg et al., 2003; Schneiders et al., 2012). Activity the prefrontal cortex of macaque monkeys has also been linked to decision making during auditory categorization tasks (Gifford III et al., 2005; Lee et al., 2009). Collectively, the ROIs automatically decoded using our machine learning approach suggest both lower-order sensory processes and higher-level cognitive functions, even during early stimulus encoding, may prime an individual for successful speech identification and subsequent behavioral actions.

## 5. CONCLUSIONS

Our findings demonstrate that categorical speech perception is supported by temporally specific neural microstates—quasi-stable patterns of distributed brain activity that reflect coordinated large-scale processing over brief time intervals (Khanna et al., 2015; Michel & Koenig, 2018). In this framework, the numbered microstate segments reflect distinct, sequential processing stages during speech categorization rather than arbitrary time bins: early post-stimulus states likely index sensory–perceptual encoding of acoustic–phonetic features (overlapping canonical N1/P2 latencies), whereas later states may reflect higher-order evaluation, decision formation, and post-perceptual/response-related processes that unfold after initial stimulus encoding. Importantly, our results show that category information is not uniformly represented across the epoch; instead, decoding peaks within a specific microstate occurring ∼197–258 ms, consistent with a temporally discrete stage at which neural representations most strongly differentiate prototypical from ambiguous tokens.

These findings align with and extend our prior microstate-based decoding work using Bayesian nonparametrics at the sensor level, which demonstrated that microstate dynamics track the speed of phonetic decisions (reaction time) during speech categorization (Al-Fahad et al., 2021). The present study advances this framework by applying microstate inference to source-reconstructed EEG, enabling identification of cortical generators and linking microstate-specific neural activity to individual differences in categorical gradiency, indexed by listeners’ identification slopes. By integrating Bayesian nonparametric state modeling, machine learning, and brain–behavior analyses, this work provides a data-driven account of how temporally discrete neural states organize the spatiotemporal dynamics underlying categorical speech perception.

One limitation of this work is that all EEG data were recorded from young adults with normal hearing ability, limits the generalizability of the findings. Categorical speech perception varies with age, hearing status, language experience, and neurological condition. Extending this approach to older adults, hearing-impaired listeners, bilingual populations, or clinical groups (e.g., dyslexia or aphasia) will be necessary to determine the broader applicability of microstate-based neural decoding.

## 6. ACKNOWLEDGMENTS

Requests for data and materials should be directed to G.M.B. This work was supported by the National Institutes of Health (NIH/NIDCD R01DC016267) and department of Electrical and Computer Engineering at the University of Memphis.

